# Doxycycline inhibits both apicoplast and mitochondrial translation in apicomplexan parasites

**DOI:** 10.64898/2026.03.11.710986

**Authors:** Michaela S. Bulloch, Emily M. Crisafulli, Jenni A. Hayward, SaiShyam Ramesh, Andrew E. Maclean, Linden Muellner-Wong, Shuai Nie, David A. Stroud, Lilach Sheiner, Alexander G. Maier, Giel G van Dooren, Stuart A. Ralph

**Affiliations:** Department of Biochemistry & Pharmacology, Bio21 Molecular Science and Biotechnology Institute, The University of Melbourne, 3010, Australia; Research School of Biology, Australian National University, Canberra, ACT, Australia; Centre for Parasitology, University of Glasgow, Glasgow, Scotland, UK; Melbourne Mass Spectrometry and Proteomics Facility, Bio21 Molecular Science and Biotechnology Institute, The University of Melbourne, 3010, Australia; Murdoch Children’s Research Institute, Royal Children’s Hospital, Melbourne, Victoria, 3052, Australia; Victorian Clinical Genetics Services, Parkville, Victoria, 3052, Australia

## Abstract

Doxycycline is a tetracycline-class antibiotic used for malarial prophylaxis and as an occasional partner drug in malaria treatment. Doxycycline’s antimalarial mechanism of action is attributed to inhibiting the prokaryotic 70S ribosomes of the apicoplast, a non-photosynthetic plastid present in most apicomplexan species including *Plasmodium* and *Toxoplasma*. At lower concentrations (<5 μM) doxycycline exhibits a delayed death phenotype, typical of inhibitors of apicoplast housekeeping processes. Unlike other delayed death drugs (e.g. clindamycin), doxycycline has rapid schizonticidal activity at higher concentrations (>10 μM) via an unknown and apicoplast-independent mechanism. In other eukaryotes, and plausibly in *Plasmodium* and *Toxoplasma*, doxycycline inhibits mitochondrial 70S ribosomes. Here we use a mass spectrometry approach to assess steady state and turnover for apicoplast DNA encoded proteins in *Plasmodium falciparum*. We directly show that these proteins decrease in both abundance and synthesis following treatment with doxycycline and clindamycin. High concentrations of doxycycline, but not clindamycin, also reduced the abundance of a mitochondrial-encoded protein. Proteins encoded by the mitochondrial genome are required for the formation of complexes III and IV in the electron transport chain. We show a reduction in abundance and activity of complex IV following doxycycline treatment in *Toxoplasma gondii*. High doxycycline concentrations also disrupt oxidative phosphorylation in both *P. falciparum* and *T. gondii*, likely as a consequence of impaired complex III and IV formation. For the first time we directly measure apicoplast translation inhibition and characterise doxycycline as a mitochondrial translation inhibitor of *P. falciparum* and *T. gondii*.

## Introduction

The apicomplexan parasites *Plasmodium falciparum* and *Toxoplasma gondii* are responsible for significant mortality and morbidity as the virulent causes of malaria and toxoplasmosis. A shared feature of these parasites is the possession of two organelles derived from endosymbiotic events: a plastid, known as the apicoplast, and the mitochondrion. These organelles are semi-autonomous, with their own prokaryotic-like genomes, proteins, and metabolic processes. Since acquisition of these organelles, the vast majority of their genes have been translocated to the nuclear genome, with the burden and control of their transcription and translation falling to the cytosolic eukaryotic machinery. N-terminal localisation sequences are predicted to direct several hundred of these nuclear DNA encoded proteins to the apicoplast and mitochondria (Boucher et al., 2018; van Esveld et al., 2021). The circular apicoplast genomes of both *P. falciparum* and *T. gondii* are ∼35 kb, with only 30 and 28 predicted protein-coding genes respectively. The apicoplast predominantly encodes genes for transcription and translation such as RNA polymerase subunits, ribosomal RNAs, transfer RNAs, and ribosomal proteins. The mitochondrial genomes are even more reduced, with only fragmented rRNA genes, and three protein-coding genes, each for subunits of electron transport chain (ETC) complexes III and IV, (Namasivayam et al., 2021; Vaidya et al., 1989; Wilson et al., 1996; Wilson et al., 1991). Despite their reduced size, the evolutionary retention of these organellar genomes suggests they are essential and, in the case of the apicoplast at least, their transcription and translation have been directly shown to be essential for parasite viability.

The presence of prokaryotic derived organelles offers many opportunities to exploit parasite specific biology for drug discovery and development, particularly organellar translation. Several clinically relevant antibiotics such as doxycycline, clindamycin and azithromycin have their anti-malarial activity attributed to inhibition of apicoplast ribosomes (Camps et al., 2002; Dahl et al., 2006; Goodman et al., 2007). The inability to reliably detect apicoplast-encoded proteins has meant that the direct measure of apicoplast translation has been challenging (Chaubey et al., 2005). Instead, the mechanism of action is inferred from a slow acting phenotype known as delayed death, where parasites exposed to apicoplast inhibitors only succumb to the lethal effects in the following cycle (Dahl & Rosenthal, 2007; Dahl et al., 2006; Fichera et al., 1995; Goodman et al., 2007; He et al., 2001). In the case of *Plasmodium* delayed death, these inhibitor-exposed plastids are metabolically inactive during the second cycle, defective in apicoplast growth, isopentenyl pyrophosphate (IPP) synthesis and protein import (Yeh & DeRisi, 2011). In contrast, so far there are no proposed mitochondrial translation inhibitors for *Toxoplasma* or *Plasmodium*, though the mitochondrion itself is the target of the clinically used antimalarial atovaquone, which inhibits the ETC (Goodman et al., 2017; Painter et al., 2007).

Doxycycline is a tetracycline-class antibiotic currently used for malarial prophylaxis and partnered with quinine is the recommended treatment for chloroquine resistant malaria. It is also occasionally used for treatment of toxoplasmosis. The known mechanism of action of doxycycline in bacteria is as a translation inhibitor, by binding to the 30S ribosomal subunit of 70S prokaryotic ribosomes, which blocks charged tRNA from binding to the mRNA-ribosomal complex (Chopra & Roberts, 2001). In both *Plasmodium* and *Toxoplasma*, doxycycline induces a “delayed death” phenotype that is consistent with impaired apicoplast maintenance which is rescuable by supplementations with the essential isoprenoid metabolite IPP (Dahl & Rosenthal, 2007; Yeh & DeRisi, 2011). Unlike other delayed death drugs, doxycycline administered at slightly higher concentrations is schizonticidal in the first cycle, suggesting an additional apicoplast-independent mode of action (Dahl et al., 2006). A likely secondary target for doxycycline is mitochondrial translation, with doxycycline inhibiting mitochondrial translation in other eukaryotes (Moullan et al., 2015) and evidence of other tetracyclines disrupting *P. falciparum* mitochondrial ETC function (Kiatfuengfoo et al., 1989; Prapunwattana et al., 1988).

We investigated whether mitochondrial translation is a secondary target of doxycycline in blood stage infections of *P. falciparum* and tachyzoite stages of *T. gondii*. Liquid chromatography-mass spectrometry (LC-MS) was used to detect and measure protein abundance following drug treatments. Both the steady state and nascently synthesised protein levels were assessed, using unlabelled and heavy amino acid labelled culture approaches. To test the effect of doxycycline on mitochondrial function, the efficiency of the ETC was measured using Seahorse metabolic flux assays and by assaying abundance and activity of the mitochondrial complexes containing protein subunits from mitochondria-encoded DNA. The data generated here has verified apicoplast translation as the target of both clindamycin and doxycycline and reveals the mitochondrion as a secondary target of doxycycline at higher doses, making it the first known mitochondrial translation inhibitor for both *P. falciparum* and *T. gondii*.

## Results

### Doxycycline does not inhibit cytosolic translation in *P. falciparum*

Like many other apicoplast inhibitors, doxycycline causes a delayed death effect in *P. falciparum* that is rescuable by IPP (Dahl et al., 2006; Uddin et al., 2018; Yeh & DeRisi, 2011) (**Figure 1**). However, compared to other antibiotics- such as clindamycin- the window between the concentrations required to elicit a delayed death versus an immediate effect is small, such that clinically relevant concentrations might produce first cycle killing. Light microscopic analysis of Giemsa-stained blood smears confirms a first-cycle morphological change in parasites treated with higher concentrations of doxycycline, appearing shrunken and condensed (a morphology known as pyknosis) within the first 16–20 hrs of treatment (**Figure 1A**). Concordant with previous reports where doxycycline has a 96 hr IC₅₀ of 0.3 to 2 µM and a 48 hr IC₅₀ of 3 to 13 µM and, we measured doxycycline to have a delayed death IC₅₀ of 963 nM, around ∼18-fold lower than its IC₅₀ of 17.0 μM at 48 hrs post treatment (Dahl et al., 2006; Uddin et al., 2018; Yeh & DeRisi, 2011) (**Figure 1B**). These data contrast with those for clindamycin, where the first cycle IC₅₀ is >100 μM – over 76,000-fold greater than its delayed death IC₅₀ of 1.31 nM (**Figure 1B**). Taken together, this suggests that doxycycline inhibits an additional target, causing an immediate killing effect not seen for other delayed death drugs at concentrations achieved *in vivo* (Newton et al., 2005).

**Figure 1.**
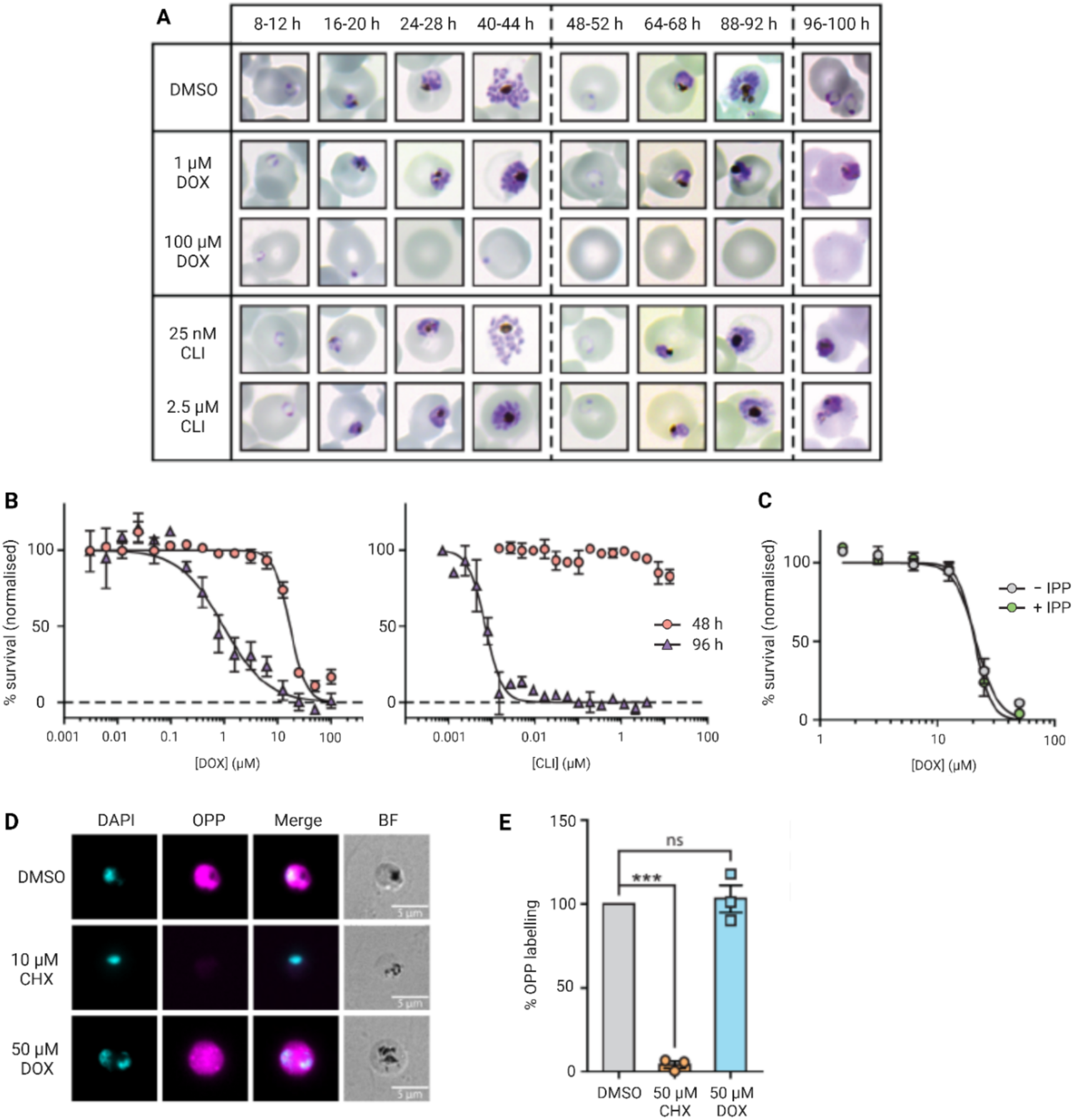
Doxycycline causes first cycle killing in *Plasmodium falciparum* that is not rescuable by isopentenyl pyrophosphate. **A)** 3D7 *P. falciparum* ring stage parasites were treated with DMSO, 1 or 100 µM doxycycline (DOX), or 25 nM or 2.5 µM clindamycin (CLI). Analysis by light microscopic analysis of Giemsa-stained blood smears over three rounds of asexual replication reveals condensed and shrunken parasites within the first replicative cycle of parasites treated with high DOX concentrations. Representative images are presented (n = 4). **B)** The first and second cycle IC50 of DOX and CLI was measured by drug treating ring stage parasites and harvesting them after 48- (orange circles) and 96- hrs (purple triangles). **C)** To probe whether the first cycle killing by DOX is mediated through an apicoplast target, DOX treated parasites were cultured in the presence (green) and absence (grey) of 200 µM isopentenyl pyrophosphate (IPP), an essential apicoplast metabolite, and harvested after 48 hrs. **B and C)** Parasites were lysed and incubated with SYBR Green to determine the relative survival. Data were normalised to a kill control (100 µM chloroquine) and presented as mean ± SEM (n = 3). **D)** Cytoplasmic translation was measured in trophozoite stage parasites which were treated with DMSO, 10 μM cycloheximide (CHX; cytosolic translation inhibitor), or 50 μM DOX for 2.5 hrs. O-propargyl-puromycin (OPP; magenta) was spiked into cultures for 30 mins, before samples were fixed and permeabilised. Parasites were incubated in the ClickiT® reaction cocktail for 30 mins and nuclei stained with 300 nM DAPI (cyan) before imaging by fluorescence microscopy. Representative data are presented (n ≥ 3). **E)** To quantify OPP fluorescence as a proxy for cytoplasmic translation, trophozoite stage parasites were treated with DMSO (grey), CHX (orange circles), or DOX (blue squares) for 2 hrs before OPP was added for another hour. Parasites were then fixed and permeabilised. Samples were treated with a CuAAC reaction cocktail for 1 hr, before staining with 1.25 μg/mL propidium iodide and quantitation by flow cytometry with the normalised mean ± SEM presented (n = 3).

There are conflicting conclusions in the literature as to where precisely this secondary target lies. A clear indication of an apicoplast target is rescue by supplementing culture media with IPP, which along with CoA are the essential products of the apicoplast for asexual erythrocytic stages, however two studies have produced differing results as to whether first cycle killing can be rescued by IPP (Okada et al., 2020; Yeh & DeRisi, 2011). To reexamine this, we supplemented ring stage parasites with 200 μM IPP and treated them with doxycycline for 48 hrs. In repeated experiments across two laboratories, we saw no significant change in the IC₅₀ of doxycycline-treated parasites in the presence or absence of IPP (**Figure 1C**)- a finding concordant with early reports (Yeh & DeRisi, 2011), although a careful analysis in multiple strains by Okada and colleagues does show a small but reproducible shift in 48-hr sensitivity under IPP rescue. Our data are consistent with an additional non-apicoplast target for doxycycline during the first cycle.

We sought to determine whether higher doxycycline concentrations may be lethal in the first cycle as a result of cytosolic translation inhibition. Active cytosolic translation was detected with a modified form of puromycin, O-propargyl-puromycin (OPP), which binds to and causes premature termination of nascent peptides. OPP is conjugated to a fluorophore, allowing for the visualisation and quantification of translation by fluorescence microscopy and flow cytometry (Liu et al., 2012; Signer et al., 2014). To determine whether we could detect organellar protein synthesis by visualising OPP-labelling, we treated parasites with 10 μM cycloheximide (a cytosolic translation inhibitor) or 50 μM doxycycline for 3 hrs. The fluorescent signal of OPP conjugated to the fluorophore was detected by fluorescence microscopy (**Figure 1D**) and quantified by flow cytometry (**Figure 1E**). While cycloheximide-treated parasites exhibited an ∼24-fold reduction in cytosolic translation, concentrations of doxycycline that elicit first cycle death had no effect on the OPP-signal, suggesting that cytosolic translation is not the lethal target of doxycycline at these concentrations. We had conjectured that blocking cytosolic translation by cycloheximide might allow visualisation of a residual weak organellar signal that had been obscured by the stronger cytosolic signal. However, no such fluorescent puncta were apparent after cycloheximide treatment, therefore, in our hands at least, this assay wasn’t useful for discerning a separate organellar (mitochondrial or apicoplast) translation activity.

### Delayed death concentrations of doxycycline and clindamycin reduce abundance of apicoplast encoded proteins

Doxycycline and clindamycin are both proposed apicoplast translation inhibitors, likely binding to the 30S and 50S ribosomal subunits respectively. To measure whether there are changes in the abundance of apicoplast DNA encoded proteins following treatment with doxycycline or clindamycin, an LC-MS approach was used. The organelle-encoded proteins of *Plasmodium* are difficult to detect using MS-based approaches in asexual blood stages (Cobbold et al., 2021; Evers et al., 2021). Apicoplast-encoded proteins have proved equally challenging, with minimal detection by LC-MS, and only one (elongation factor Tu, Pf3D7_API02900) by immunoprecipitation (Chaubey et al., 2005; Hollin et al., 2025). Therefore, to obtain as deep protein coverage as possible, strong cation exchange (SCX) fractionated samples of *P. falciparum* cell lysate were used to generate a peptide spectral library that supplemented mass spectrometry peak matching by data independent acquisition (DIA) in subsequent experiments. Biochemical techniques can also be used to enrich for organelle-specific proteins such as cell fractionation, differential permeabilization or immunoprecipitation of tagged organelles, however we favoured an untargeted approach to assess global protein changes that might additionally reveal the involvement of non-organellar pathways (Evers et al., 2021). These facilitated the detection of 1 mitochondrial DNA encoded protein and 10 apicoplast DNA encoded proteins (summarised in **Table 1**). The log₂ transformed protein abundances of the organellar-encoded proteins for each replicate (prior to imputation of missing values at the limit of detection) is included in **Supplementary Tables 1-7**.

**Table 1.** Summary of *P. falciparum* apicoplast and mitochondrion DNA encoded proteins detected by LC-MS in this study.

| PlasmoDB ID | Protein name |
| --- | --- |
| Pf3D7_API01700 | Ribosomal protein S3, putative |
| Pf3D7_API01800 | Apicoplast ribosomal protein L16 |
| Pf3D7_API02700 | Apicoplast ribosomal protein S12 |
| Pf3D7_API02900 | Elongation factor Tu |
| Pf3D7_API03600 | Chaperone protein ClpM |
| Pf3D7_API04000 | Hypothetical chloroplast reading frame 93 |
| Pf3D7_API04100 | 30S ribosomal protein S2, putative |
| Pf3D7_API04200 | DNA-directed RNA polymerase subunit beta, putative |
| Pf3D7_API04400 | DNA-directed RNA polymerase subunit beta, putative |
| Pf3D7_API04500 | Hypothetical protein |
| Pf3D7_MIT02300 | Cytochrome *b* |

To monitor and compare the steady state proteomic profile throughout delayed death, synchronised ring stage parasites were exposed to 1 µM doxycycline or 25 nM clindamycin (concentrations sufficient to generate delayed death but not first cycle death). Parasites were then harvested after 24 or 72 hrs, corresponding to the trophozoite stages in the first and second cycles, and proteins quantified by MS (**Figure 2**). While few significant (defined by a fold change of >1.5 in either direction, p<0.05) changes were observed in the first cycle, by the second cycle a decrease in both apicoplast-encoded and targeted proteins was detected in both drug treatments (**Figure 2**). Parasites treated with clindamycin saw an unexpected increase in abundances of numerous proteins by 72 hrs that was not observed in the doxycycline samples.

**Figure 2.**
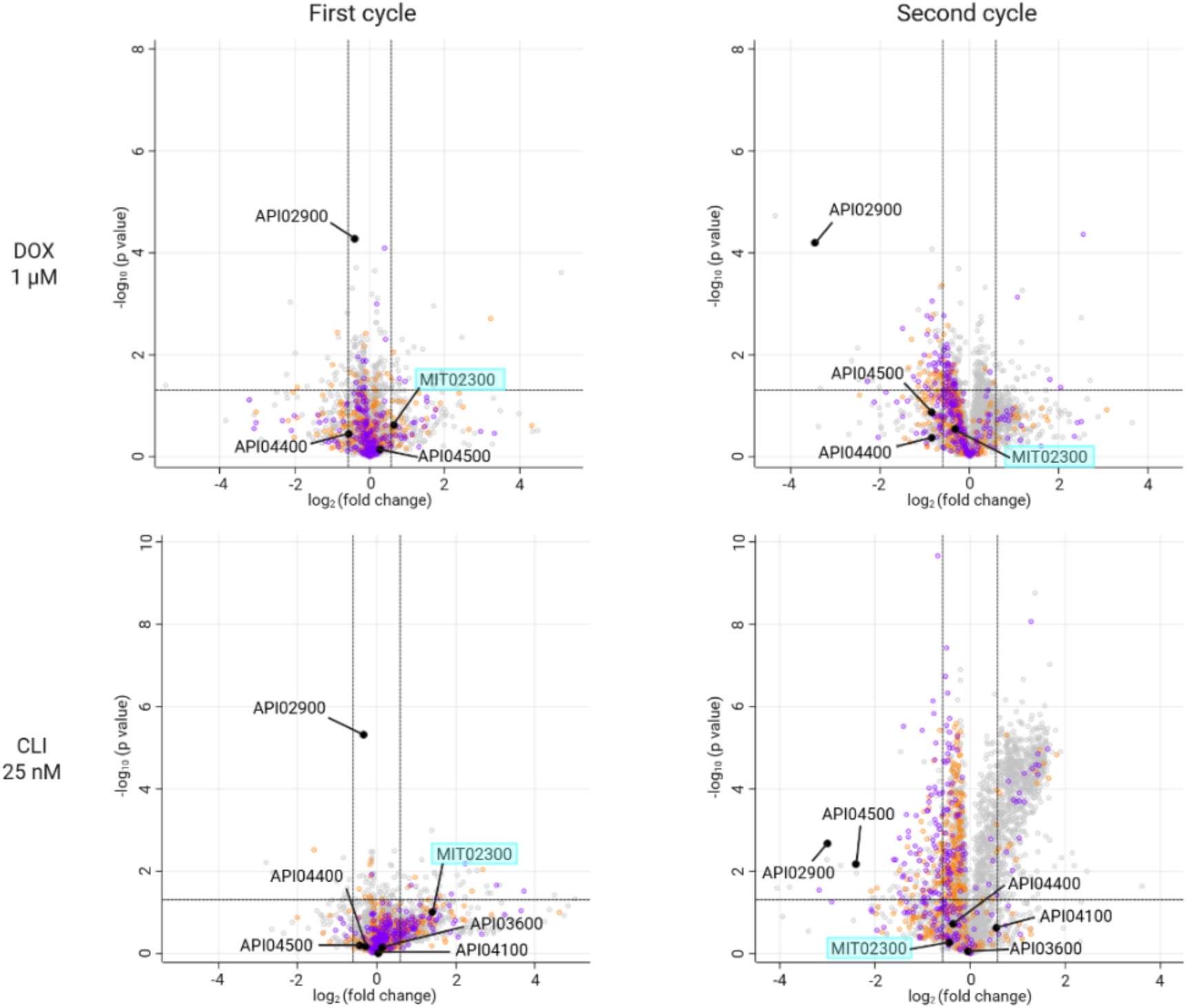
Apicoplast DNA encoded proteins are reduced in second cycle *P. falciparum* during delayed death. Ring stage parasites were treated with delayed death concentrations of doxycycline (DOX, 1 µM) or clindamycin (CLI, 25 nM) and harvested at the trophozoite stage in the first and second cycle. An LC-MS proteomics approach was used to detect and measure the changes in protein abundance following treatment, and statistical changes in protein abundance are displayed using a volcano plot. Proteins derived from either the apicoplast or mitochondrial genomes are annotated in black with the PlasmoDB IDs (gene Ids starting with API or MIT respectively). Nuclear-encoded proteins targeted to the apicoplast and mitochondria are coloured purple or orange respectively. All other proteins are light grey. A significance threshold of the p-value < 0.05 and a fold change of > 1.5 was set. For doxycycline treatments, there were 3 DMSO control and 3 drug treated replicates, while for clindamycin treatments 5 DMSO control and 5 drug treated replicates were used.

### High concentrations of doxycycline inhibit mitochondrial translation

At concentrations only around 10-fold higher than its delayed death IC₅₀, doxycycline is lethal in the first cycle by an apicoplast independent mechanism. However, for clindamycin treatments, even 1,000 x the delayed death IC₅₀ is insufficient for a first cycle phenotype (Dahl & Rosenthal, 2007). To distinguish the proteomic changes following high drug concentrations, trophozoite stage cultures were exposed to 25 x the delayed death IC₅₀ concentration of doxycycline and clindamycin (25 µM and 625 nM respectively). The dihydrofolate reductase (folate metabolism) inhibitor WR99210 was included as a non-apicoplast targeting control at the lethal concentration of 5 nM (Fidock & Wellems, 1997). After an 8-hr drug pulse, trophozoite parasites were harvested for proteomic analysis. Doxycycline and clindamycin, but not WR99210 induced a decrease in the abundance of apicoplast-encoded proteins (**Figure 3A-C**). Furthermore, 25 µM doxycycline was the only treatment to significantly reduce abundance of the only detectable mitochondrial DNA encoded protein (cytochrome *b*), plausibly due to the inhibition of mitochondrial translation (**Figure 3A**). This concentration of doxycycline also induced many non-mitochondrial and non-apicoplast protein changes, indicative of the wider cellular impact evident by the pyknotic parasites seen by Giemsa (Figure 1A).

**Figure 3.**
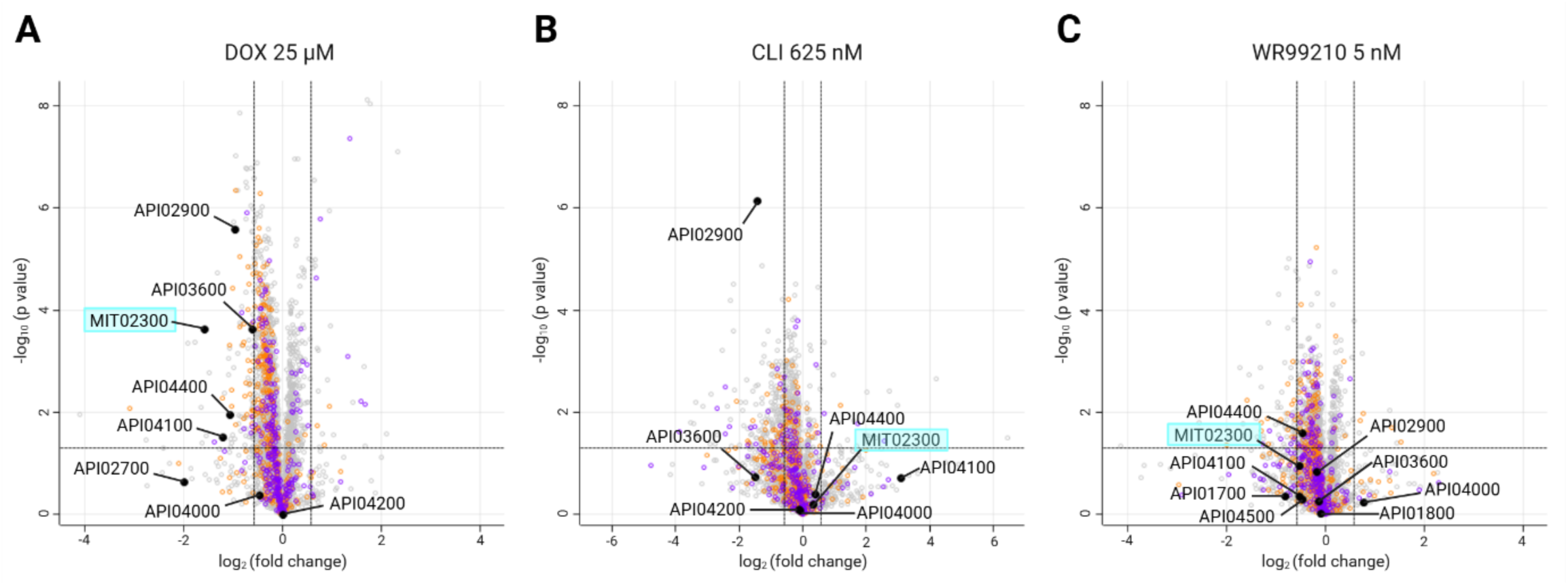
Doxycycline inhibits mitochondrial protein synthesis. Trophozoite stage *P. falciparum* were exposed to 25 x the delayed death IC₅₀ of **A)** doxycycline (DOX, 25 µM), **B)** clindamycin (CLI, 625 nM) or **C)** dihydrofolate reductase inhibitor WR99210 as a control treatment that doesn’t target protein translation (at the lethal concentration of 5 nM) for 8 hrs. Changes in protein abundance was measured by LC-MS. Of proteins derived from organellar genomes, apicoplast and mitochondrion proteins (annotated in black) decreased following doxycycline treatment, while an apicoplast but not mitochondrial protein was affected by clindamycin. No organellar-encoded protein were significantly different following WR99210 treatment. Nuclear-encoded proteins targeted to the apicoplast, and mitochondria are coloured purple or orange respectively, with all other proteins light grey. A significance threshold of the p-value < 0.05 and a fold change of > 1.5 was set. The doxycycline and clindamycin treatments each had 5 replicates, with 5 corresponding DMSO controls), while the experiment with WR had 4 DMSO and 4 drug treated replicates.

To determine whether there is a proteomic signature or universal response to organellar translation inhibitors, the proteins that were significantly increased and decreased were compared across clindamycin and doxycycline conditions. Time points where we have demonstrated organellar translation inhibition (i.e. second cycle trophozoites during delayed death, and first cycle trophozoites exposed to high concentration of doxycycline) were chosen as these are actively affected. Cross-analysis revealed that there were three proteins commonly decreased in abundance, one being an apicoplast-encoded protein (elongation factor Tu) (**Supp Figure 1A**). The four commonly increased proteins had unknown functions with the exception of a folate-biopterin transporter, a transporter type that has been implicated in anti-folate drug resistance mechanisms (**Supp Figure 1B**).

### Rapid inhibition of translation can be detected by pulse SILAC mass spectrometry

The steady state proteome is influenced not only by protein translation, but other factors such as protein stability and degradation. The medium-term persistence of extant proteins after application of an inhibitor can have the effect of diluting any detectable impact on newly-synthesised proteins. To measure new protein translation directly, pulse stable isotope labelled amino acid culturing (SILAC) was used. All amino acids used by the parasite can be derived from haemoglobin digestion except isoleucine, making this the only amino acid that must be acquired extracellularly (which during *ex vivo* cultivation is provided in the growth media) (Liu et al., 2006). Therefore, labelled isoleucine was used for the SILAC experiments (Nirmalan et al., 2004). Synchronised trophozoites cultured in media with light isotope amino acids were exposed to doxycycline (at 1 µM or 25 µM), clindamycin (25 nM) or WR99210 (5 nM) for a total of 8-hrs. After 3 hrs, the media was replaced with that containing heavy isotope isoleucine (¹³C¹⁵N-isoleucine) and exposed to drug for another 5 hrs before the parasites were harvested. Any proteins synthesised in the 5-hr window should incorporate the heavy isoleucine and be distinguishable from those with only light amino acids (**Figure 4A**) (Yang et al., 2019).

**Figure 4.**
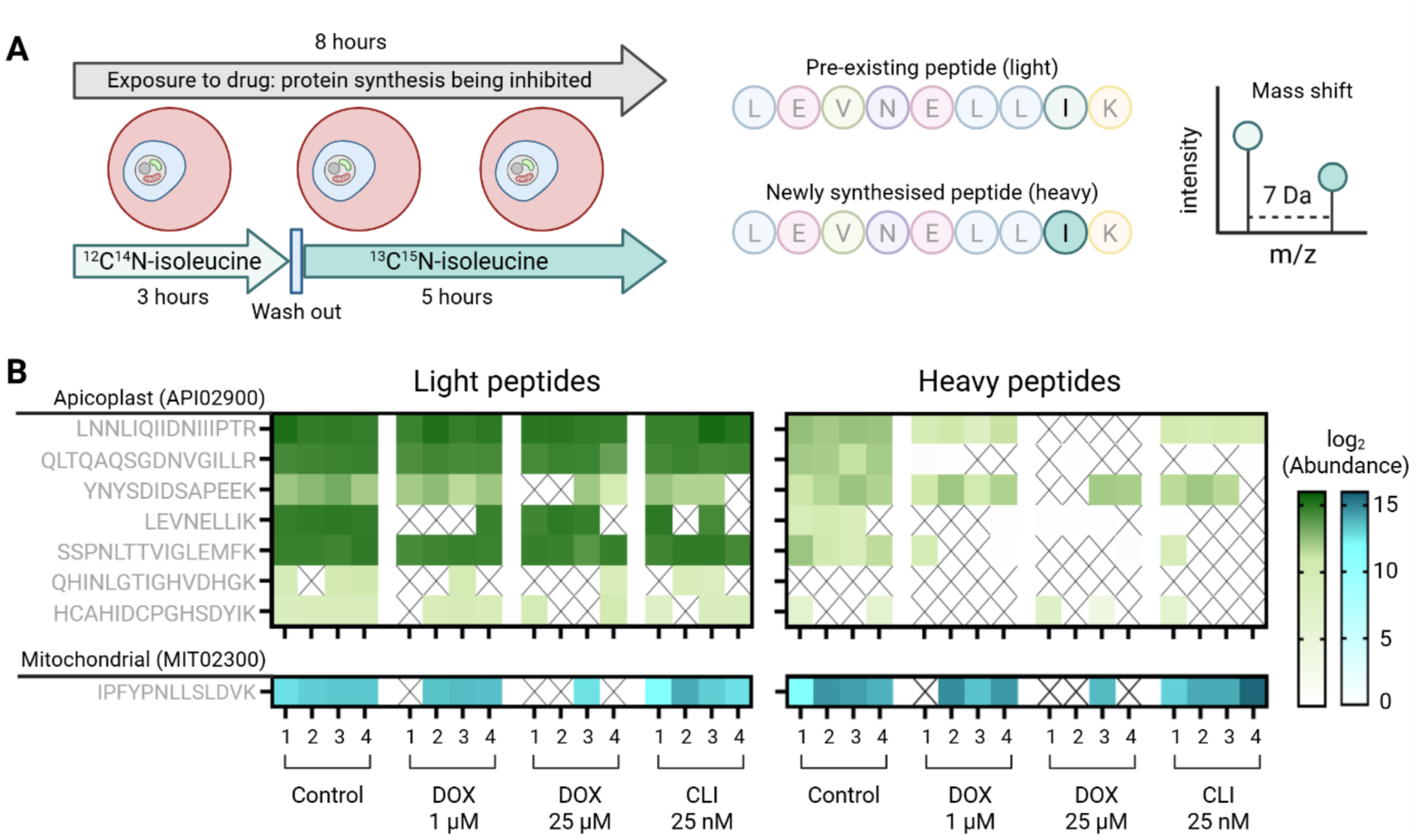
Pulse SILAC shows rapid inhibition of apicoplast translation in *P. falciparum*. **A)** Trophozoite stage *P. falciparum* were exposed to translation inhibitors for a total of 8 hrs. After the first 3 hrs, the media was exchanged from that containing light amino acids to that with heavy isoleucine (¹³C¹⁵N-isoleucine). Any newly synthesised proteins will incorporate the heavy isoleucine, which can be distinguished from those with light isoleucine by mass spectrometry. **B)** Heat map showing the log₂ abundance of the light and heavy isoleucine-containing peptides of organellar-encoded proteins. The four replicates for each treatment are shown. Only light peptides detected in at least 75% of replicates for at least one treatment are shown.

The isoleucine containing peptides of the organellar-encoded proteins were assessed. Light peptides from one apicoplast and one mitochondrial encoded-protein could be detected in each of the treatments, validating that these proteins are present in the samples prior to drug exposure. While heavy peptides (corresponding to newly translated protein) from both the apicoplast and mitochondrion encoded proteins detected in our dataset are mostly present in the control parasites, many of these are significantly decreased or missing altogether from the doxycycline and clindamycin (**Figure 4B**) but not WR99210 (**Supp Figure 2**) treatments. Together, the steady state and SILAC LC-MS data (summarised in **Table 2**) confirms apicoplast translation inhibition by both doxycycline and clindamycin, with the increased temporal sensitivity of the SILAC method revealing that inhibition occurs within the first 8 hrs of treatment. Unfortunately, only one isoleucine-containing peptide from the mitochondrial DNA encoded protein met our inclusion criterion, making it difficult to draw conclusions about inhibition of mitochondrial translation by SILAC LC-MS.

**Table 2.** Summary of LC-MS experiments and whether organellar translation inhibition could be detected using either the steady state or SILAC method. Parasites were exposed to doxycycline (DOX) at 1 µM (low) or 25 µM (high), clindamycin (CLI) at 25 nM (low) or 625 nM (high), and WR99210 (WR) at 5 nM. YES = at least one protein had a statistically significant decrease in protein abundance following treatment (p value < 0.05, fold change > 1.5); NO = no organellar-encoded proteins were found to be statistically significantly different following treatment; N/A = not assessed; N/D = not determined due to inability to detect peptides.

|  | Apicoplast translation inhibition |  | Mitochondrial translation inhibition |  |
| --- | --- | --- | --- | --- |
|  | Steady state | SILAC | Steady state | SILAC |
| Low DOX 1<sup>st</sup> cycle | NO | YES | NO | NO |
| Low DOX 2<sup>nd</sup> cycle | YES | N/A | NO | N/A |
| High DOX 1<sup>st</sup> cycle | YES | YES | YES | NO |
| Low CLI 1<sup>st</sup> cycle | NO | YES | NO | NO |
| Low CLI 2<sup>nd</sup> cycle | YES | N/A | NO | N/A |
| High CLI 1<sup>st</sup> cycle | YES | YES | NO | NO |
| WR 1<sup>st</sup> cycle | NO | NO | NO | N/D |

### Abundance of *P. falciparum* ETC proteins are minimally perturbed following drug treatments

An indirect measure of mitochondrial translation is the stability and function of the ETC complexes III and IV, which are comprised of subunits encoded on both the mitochondrial and nuclear genomes, as opposed to complexes II and V which consist entirely of nuclear encoded-proteins (Evers et al., 2021). Given that high concentrations of doxycycline reduce the abundance of the only detectible mitochondrially-encoded protein, cytochrome *b*, the changes in abundance of nuclear encoded subunits of complexes II-V were monitored by LC-MS across the different treatments. While we observed a decrease in the abundance of cytochrome *b* via LC-MS after 8 hrs doxycycline exposure, major changes to proteins in complexes III and IV were not detected during this time frame (**Supp Figure 3**).

### Doxycycline inhibits mitochondrial ETC activity of *P. falciparum* and *T. gondii*

While the ETC components of *P. falciparum* are still present following doxycycline treatment, we sought to determine whether ETC function may nonetheless be compromised. Uptake of the mitochondrial membrane potential-dependent dye MitoTracker Red was reduced by increasing concentrations of doxycycline, suggesting that ETC activity is being affected (**Supp Figure 4**). To probe the activity of the ETC complexes individually, a Seahorse metabolic flux assay was performed (Ramesh et al., 2023). These assays determine the mitochondrial oxygen consumption rate (mOCR) as a measure of ETC activity. The functionality of individual ETC complexes can be separately determined through the addition of various compounds during the assay. First, the complex III inhibitor atovaquone is added to provide a measure of complex III-dependent ETC activity. Next, N,N,N′,N′ - tetramethyl-p-phenylenediamine dihydrochloride (TMPD), a compound that donates electrons directly to cytochrome *c* by bypassing complex III, is added, providing a measure of complex IV- dependent ETC activity. Finally, the complex IV inhibitor sodium azide (NaN₃) is added to validate that TMPD addition is providing a measure of complex IV activity (**Figure 5A**).

**Figure 5.**
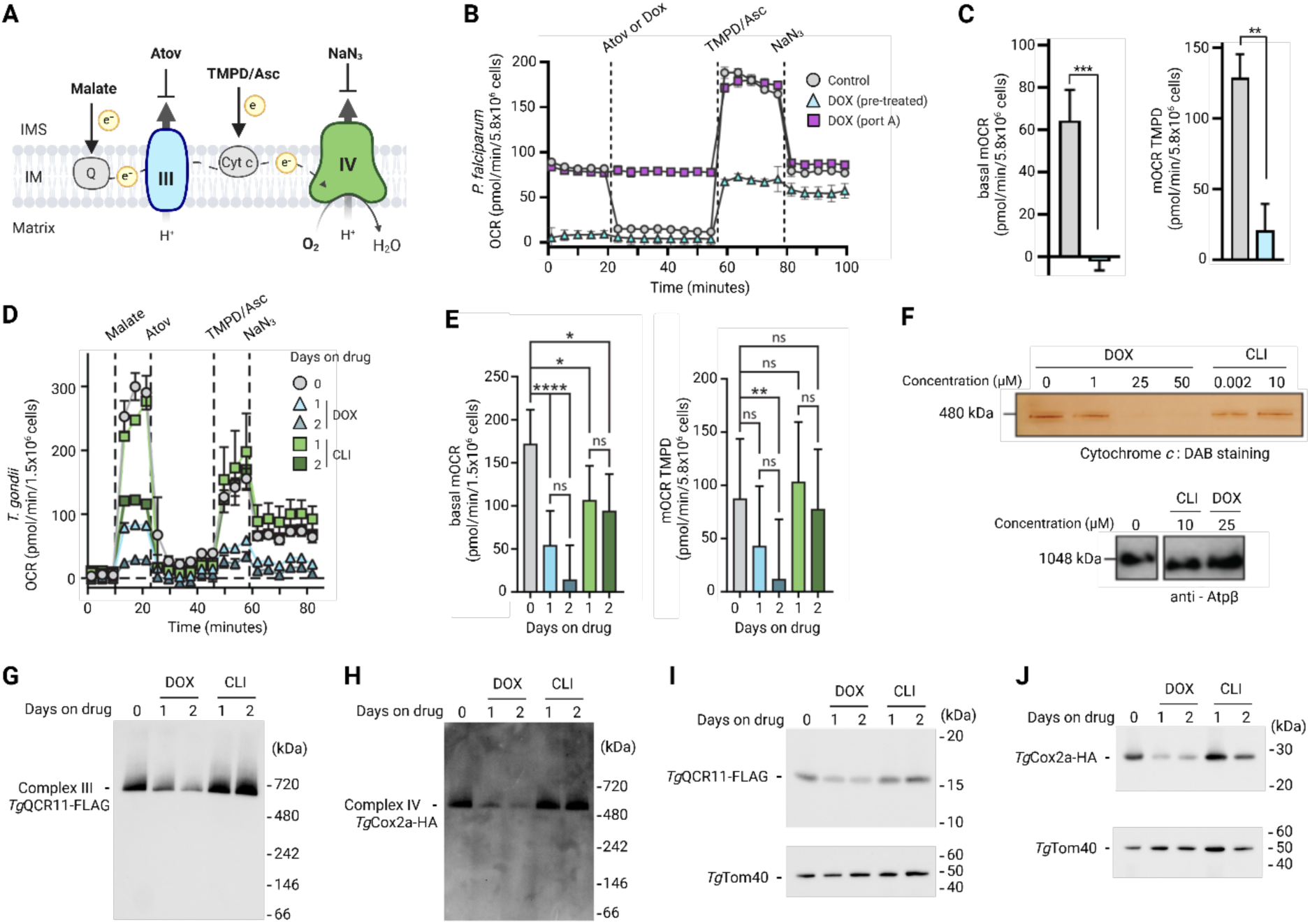
Measuring the ETC efficiency in *P. falciparum and T. gondii*. **A)** Seahorse metabolic flux assays were used to measure the activity of the electron transport chain (ETC) through determining the oxygen consumption rate (OCR). ETC complex inhibitors such as atovaquone (Atov) and NaN₃ decrease the OCR while electron donors such as malate and TMPD, reduced by ascorbate (Asc), can deliver electrons directly to ubiquinone (Q) and cytochrome *c* (Cyt *c*) respectively. **B)** Example trace of *Plasmodium falciparum* exposed to DMSO (control, grey circles) or 25 µM doxycycline either for ∼15 hrs prior to the experiment (DOX, blue triangles) or spiked in during the assay (purple squares). **C)** Pre-treatment with doxycycline severely reduced basal mOCR (initial malate-dependent OCR minus OCR after Atov injection) and TMPD-dependent mOCR (OCR following TMPD injection minus OCR after Atov injection) compared to the DMSO control. Data presented as mean ± SD (n = 3 biological replicates (2-3 technical replicates per condition)). Unpaired t-test, p <0.05 *, <0.01 **, <0.001 ***. **D)** OCR measurements of *Toxoplasma gondii* permeabilised parasites in the presence of either DMSO, 24 µM doxycycline or 10 µM clindamycin (CLI, green squares) for 1-2 days. Data presented as mean ± SD (n = 3) **E)** Basal malate-elicited and TMPD-elicited mOCR of parasites grown in the presence of DMSO, DOX, or CLI for 1–2 days. Data depict the mean ± 95% CI of the linear mixed-effects model (n = 3). ANOVA followed by Tukey’s multiple pairwise comparisons test was performed: *p ≤ 0.05; **p ≤ 0.01; ***p ≤ 0.001; ****p ≤ 0.0001. **F)** *T. gondii* D*ku*80/TATi tachyzoites were treated for 24 hrs with 1, 25 or 50 µM DOX, or 0.002 or 10 µM CLI. Whole cells we solubilised in 2% (w/v) DDM and the soluble material loaded for clear or blue native PAGE analysis of mitochondrial protein complexes. Complex IV activity was analysed using an in-gel cytochrome *c*: 3,3’- diaminobenzidine (DAB) stain, and complex V assembly by Western blot (representative plot shown), probing with rabbit anti-Atpβ. **G-J**) Western blots of proteins extracted from *Tg*Cox2a-HA/*Tg*QCR11-FLAG parasites cultured in the presence of DMSO, 25 µM DOX or 10 µM CLI for 1-2 days, separated by (**G-H**) BN-PAGE or (**I-J**) SDS-PAGE, and probed with anti-FLAG antibodies to detect the complex III protein *Tg*QCR11-FLAG, anti-HA antibodies to detect the complex IV protein *Tg*Cox2a-HA, or anti-*Tg*Tom40 antibodies as a loading control. Data are representative of two independent experiments.

To determine the OCR changes in *P. falciparum* following doxycycline treatment, ring-stage parasites were pre-treated with 25 µM doxycycline and then harvested ∼15 hrs later, during the trophozoite stage. The parasites were then released from the host red blood cell and the plasma membrane permeabilised with digitonin to enable the ETC substrate malate to enter the mitochondrion and stimulate ETC activity to provide a measure of the basal OCR. Pre-treatment with doxycycline, but not injection during the assay, perturbs OCR, indicating that doxycycline itself does not directly inhibit electron transport chain activity (**Figure 5B**). This is consistent with doxycycline having an indirect impact by blocking translation of mitochondrial DNA encoded subunits for complexes III and IV the electron transport chain. This longer doxycycline exposure resulted in a reduction to negligible basal OCR activity and with only a minor increase in OCR following TMPD addition, suggesting complex IV activity is compromised (**Figure 5C**).

Antibiotic treatment of the related apicomplexan parasite *T. gondii* with tetracyclines also appears to have an additional apicoplast-independent target, which has been suggested to be mitochondrial ribosomes (Beckers et al., 1995; Camps et al., 2002). Like *P. falciparum*, complexes III and IV of the *T. gondii* ETC contain mitochondrial DNA-encoded subunits. Therefore, to assess whether doxycycline also impairs ETC activity of *T. gondii*, tachyzoites stage parasites were treated for 1-2 days with both low and high concentrations of doxycycline or clindamycin (**Figure 5D**). We observed a ∼70% decrease in malate-dependent OCR after one day treatment with doxycycline and a >90% decrease after two days, which mirror our findings in *P. falciparum*. Surprisingly, we did also see an impairment of malate-dependent OCR upon clindamycin treatment, resulting in a ∼40-50% decrease (**Figure 5E**). The reduction in OCR was maintained following the addition of TMPD in doxycycline treated parasites, but were restored in clindamycin treatments, demonstrating that clindamycin has no effect on complex IV of the ETC. This was recapitulated in an in-gel complex IV activity assay, where higher concentrations of doxycycline, but not clindamycin, reduced complex IV activity whilst maintaining assembly of complex V (**Figure 5F**). The overall stability of complexes III and IV was assessed using a combination of BN-PAGE and SDS-PAGE western blots. To enable the simultaneous detection of both complexes, we introduced a HA tag into the gene encoding the complex IV protein *Tg*Cox2a in an existing parasite line in which the gene encoding the complex III protein *Tg*QCR11 was FLAG tagged (**Supp Figure 5**; (Hayward et al., 2021)). Doxycycline but not clindamycin treatment resulted in a decrease in the abundance of complexes III and IV as seen by BN-PAGE (**Figure 5G** and **5H**), as well as a reduction in the individual subunits *Tg*QCR11 and *Tg*Cox2a by SDS-PAGE (**Figure 5I** and **5J**), as would be expected if a core subunit of the complex (like mitochondrial DNA-encoded cytochrome *b*) is missing or depleted.

## Discussion

The apicoplast has long been considered a clinically significant drug target in medically important apicomplexans such as *Plasmodium* and *Toxoplasma*. Understanding the mechanism of action for any drug is important for anticipating off-target effects, safety, efficacy, potential resistance mechanisms and interference with other drugs. These concerns are especially crucial in the context of anti-malarials, which due to a long history of emergence of parasite resistance, are administered in cocktails of drug combinations. The challenges in detecting plastid-encoded proteins has made the characterisation of apicoplast translation inhibitors previously untenable. Here, we demonstrate the anti-apicoplast mechanism of action for both clindamycin and doxycycline, two clinically used anti-malarials. Additionally, we have elucidated mitochondrial translation as a secondary target for doxycycline in these parasites.

Before the discovery of the apicoplast, the anti-malarial action of many anti-microbials was attributed to the mitochondrion given its bacterial origin. The subsequent revelation of an apicoplast with its own bacterial-type ribosomes led to these being demonstrated as the primary target of doxycycline in apicomplexans (Dahl et al., 2006; Yeh & DeRisi, 2011). However, unlike most other delayed death drugs, doxycycline and other tetracyclines have a pharmacologically important secondary target, the identity of which has been debated for decades. Given their widely accepted mechanism of action against 30S ribosomal subunits, the mitochondrion was naturally suspected. Previous studies have shown that tetracycline treatments decrease activity of dihydroorotate dehydrogenase, an electron donor in the ETC, along with apicoplast and mitochondrial transcript abundance (Lin et al., 2002; Prapunwattana et al., 1988). These reports are consistent with our observations that high doses of doxycycline inhibit organellar translation and perturbs the ETC. More recently, exogenous iron has been shown to rescue *P. falciparum* parasites from high concentration doxycycline treatments in the first, but not second cycle, which has been attributed to inhibition of an unspecified metal-dependent pathway (Okada et al., 2020). The existence of a non-translation apicoplast target is further supported by the observation that apicoplast morphology in the first cycle can also be perturbed by doxycycline (Okada et al., 2020).

The mitochondrial target for doxycycline may have some clinically relevant impact on other drugs inhibiting mitochondrial functions. Limited drug interaction data are available for these pairs, but one *in vitro* study reported a mildly synergistic effect of the atovaquone-doxycycline combination (Yeo et al., 1997). A combination of atovaquone and doxycycline also showed some improvement over either drug alone in human malaria cure rates, although it was unclear if any interaction above additivity was present (Looareesuwan et al., 1996). Despite being widely used for malaria prophylaxis and treatment since the 1990s, reports of genuine failure of doxycycline treatment or prophylaxis are sparse, and the establishment of substantially resistant parasites is yet to be reported (see (Home et al., 2025) for a discussion). Doxycycline resistance in other organisms such as bacteria is certainly possible, and is conferred through rRNA binding site mutations, ribosomal protection proteins, mutations in and copy number variations in efflux pumps and enzymatic inactivation (Chopra & Roberts, 2001; Grossman, 2016). While these resistance mechanisms can spread between bacteria through mobile genetic elements, this phenotype is harder to acquire in the malaria parasite (Partridge et al., 2018).

We have established an assay that allows for the detection and quantification of apicoplast and mitochondrial DNA-encoded proteins, both newly translated and steady state levels, so aspects of organellar biology previously only predicted could now be investigated. Importantly, these assays demonstrate direct inhibition of apicoplast translation that reconfirms this as the primary target of doxycycline. However, our data also for the first time establishes pharmacological inhibition of mitochondrial translation in *Plasmodium* and reveal this as a likely contributor to first-cycle killing by doxycycline. The potency in this case is low, so does not yet provide a compelling example for mitochondrial translation as an attractive drug target, but the observations certainly point at mitochondrial translation as a potential relevant targetable pathway. Now that we have evidence for mitochondrial translation as inhibitable, we can leverage this for drug development, with recent structural work on the mitochondrial ribosomes opening opportunities for structure-based inhibitor design (Shikha et al., 2025; Wang et al., 2024).

## Materials and methods

### Plasmodium falciparum culture

Asexual blood stage 3D7 *P. falciparum* was continuously cultured as previously described in O positive human red blood cells (supplied by the Australian Red Cross Lifeblood) at ∼4% haematocrit in complete culture media (CCM) (RPMI-1640 (Thermo Scientific), 14 mM sodium bicarbonate, 25 mM HEPES, 0.37 mM hypoxanthine and 40 µg/mL gentamycin (Thermo Scientific)) (Trager & Jensen, 1976). Infected RBCs (iRBCs) were maintained at 37°C in an atmosphere of 1% O₂, 5% CO₂, 94% N₂.

In experiments using SILAC media, cultures were started on SILAC media with unlabelled isoleucine (light media) in the cycle prior to drug treatment. Drug treatment was given for a total of 8 hrs, the first 3 while parasites were fed light media, which was replaced with media containing ¹³C₆¹⁵N₁ isoleucine (Cambridge Isotopes) (heavy media) for the remaining 5 hrs, which is the minimum required time for maximum heavy isoleucine incorporation (Yang et al., 2019).

### *P. falciparum* synchronisation

For tight synchrony of parasite age, a Percoll® (Sigma-Aldrich) enrichment of schizont stage parasites was performed prior to a sorbitol synchronisation. Enrichment of RBCs infected with schizont stage parasites was performed as previously described (Rivadeneira et al., 1983). Cultures were loaded onto a cushion of 65% (v/v) Percoll® in PBS and centrifuged. Schizont iRBCs at the interface of the CCM and Percoll layers were collected and returned to culture on an orbital shaker at 37 °C to maximise merozoite re-invasion. After 4 hrs, newly invaded ring-stage parasites were synchronised with 5% (w/v) D-sorbitol to lyse any RBCs infected with schizont stages. The intact RBCs were pelleted and returned to culture to produce a ring-stage culture synchronised to a 4-hr window.

### *Plasmodium falciparum* drug sensitivity assays

Drug treatments of *P. falciparum* were initially assessed using light microscopy. Ring stage parasites synchronised to a 4-hr window were incubated with either dimethyl sulfoxide (DMSO), 1 or 100 µM doxycycline (Sigma Aldrich, D3447), and 25 nM or 2.5 µM clindamycin (Sigma Aldrich, C5269). Blood smears were taken over three progressive cycles, stained with 10% (v/v) Giemsa and visualised by light microscopy under a 100x oil immersion objective.

For drug survival assays, tightly synchronised ring stage parasites were seeded into V-bottom 96-well plates and incubated with dose gradients of doxycycline or clindamycin for either 48 or 96 hrs in total. Wells containing parasites in the absence of drug were used as positive controls, along with kill controls where 100 µM chloroquine (Sigma Aldrich, C6628) was added (at 0 or 72 hrs for 48- and 96-hr assays respectively). In experiments with isopentenyl isoprenoid (IPP) (ISOPRENOIDS, IPP001) supplementation, 200 µM IPP was included for the duration of the assay. Technical triplicates were made for each biological replicate.

Following drug incubation, iRBCs were incubated with lysis buffer (20 mM TRIS (pH 7.5), 5 mM EDTA, 0.008% (w/v) saponin, 0.08% (v/v) Triton X-100) and SYBR Green I (Thermo Fisher Scientific) for 1 hr at 4 °C (Smilkstein et al., 2004). The fluorescent signal was detected using a FLUOstar Omega plate reader (BMG Labtech). Survival was calculated by subtracting background (chloroquine treated) signal and normalising data to the untreated control.

### O-propargyl-puromycin (OPP) assay for measuring protein translation

To visualise protein translation by fluorescence microscopy, synchronised trophozoite parasites were treated with either DMSO, 10 µM chloroheximide (Sigma Aldrich, C7698), or 50 µM doxycycline for 2.5 hrs. 20 µM O-propargyl-puromycin (OPP; Thermo Fisher Scientific) was then spiked into the cultures which were incubated for a further 30 mins. Control samples lacking OPP were concurrently prepared. Samples were fixed to glass coverslips and permeabilised with 0.1% (v/v) Triton X-100. Permeabilised parasites were subjected to a click reaction using the Click-iT™ Plus OPP Protein Synthesis Assay Kit (Thermo Fisher Scientific). Parasites were incubated in the Click-iT™ reaction cocktail (Click-iT™ OPP Reaction Buffer, 2% (v/v) copper protectant, 0.25% (v/v) Alexa Fluor™ 647 picolyl azide, and Click-iT™ Reaction Buffer Additive) for 30 min and protected from light. Control samples lacking Alexa Fluor™ 647 picolyl azide were concurrently prepared. Parasites were washed once with the Click-iT™ Reaction Rinse Buffer and stained with 300 nM DAPI. Coverslips were mounted onto glass slides for imaging. Data were processed using ImageJ (Version 1.53j) (Schindelin et al., 2012).

To quantify the OPP signal by flow cytometry, parasites were synchronised with sorbitol and at the trophozoite stage were added to V-bottom 96-well plates and treated with either DMSO, or dose gradients of cycloheximide or doxycycline for 2 hrs. 4 µM OPP was then spiked into the cultures and parasites were incubated for 1 hr. Control samples lacking OPP were concurrently prepared. Technical replicates were prepared in duplicate for each biological repeat. Following OPP incubation, parasites were washed in PBS and fixed in 4% (w/v) paraformaldehyde (PFA) and 0.02% glutaraldehyde for 20 mins. Parasites were then permeabilised with 0.05% (v/v) Triton X-100 (in PBS with 3% (v/v) serum) and then pelleted and resuspended in a copper(I)-catalysed azide-alkyne cycloaddition (CuAAC) reaction cocktail (100 µM CuSO4, 500 µM tris hydroxypropyltriazolylmethylamine (THPTA), 5 mM sodium ascorbate, and 100 µM Alexa Fluor™ 488 picolyl azide (Thermo Fisher Scientific) in PBS), and incubated at 37 °C, protected from light, for 1 hr. Control samples lacking Alexa Fluor™ 488 picolyl azide were concurrently prepared. Parasites were washed before a 5 min incubation with propidium iodide (PI; 1.25 µg/mL PI (Invitrogen) in PBS with 3% serum). Samples were diluted 1:4 with PBS (with 3% (v/v) serum) into a fresh V-bottom 96-well plate. Data were collected by flow cytometry (BD FACSCanto™ II; 488 nm laser with FITC and PerCP-Cy5.5 filters). Analysis was performed using FlowJo (Version 10.8) to determine parasitemia and identify the proportion of parasites exhibiting OPP mediated fluorescence. Background signal from the wells lacking Alexa Fluor™ 488 picolyl azide and OPP was removed before data were normalised to parasitemia and then to the untreated control.

### Drug regimens and harvesting for proteomics experiments

*P. falciparum* cultures tightly synchronised to a 4-hr window were treated with either DMSO, doxycycline (1 µM or 25 µM), clindamycin (25 nM or 625 nM) or WR99210 (Jacobus Pharmaceuticals, 5 nM). Drug pulses of 8, 24 and 72- hrs were applied, with media replaced daily for the 72-hr time point.

Parasites were harvested using 0.03% (w/v) saponin in PBS and the parasite lysate was washed in ice cold PBS with cOmplete™ mini EDTA-free protease inhibitor (Roche). The supernatant was entirely removed, and pellets were snap frozen in liquid nitrogen to be stored at -80 °C. Parasite pellets were resuspended in solubilisation buffer (5% (w/v) SDS, 50 mM triethylammonium bicarbonate (TEAB), benzonase) and incubated at room temperature for 10 mins, before sonication in a water bath for 15 mins at 30 °C. Samples were centrifuged and the supernatant collected.

### Alkylation and reduction

The total protein extracted was quantified with a bicinchoninic acid (BCA) assay (Thermo Fisher). Protein samples were alkylated with 40 mM 2-chloroacetamide (Sigma-Aldrich) and reduced with 10 mM TCEP. After vortexing briefly, samples were incubated at 99 °C with shaking for 5 mins. Orthophosphoric acid was added for a final concentration of 1.2% (v/v) and samples were vortexed again.

### S-Trap™ and trypsin digest

S-Trap binding/washing buffer (90% (v/v) methanol, 100 mM TEAB, pH 7.1) was added to the protein sample solution at a ratio of 7:1 (v/v) and then transferred to S-Trap™ mini columns (ProtiFi) (Kulak et al., 2014). The bound proteins were washed with S-Trap buffer before their overnight digestion (trypsin solubilized in 50 mM acetic acid, diluted in 50 mM TEAB for a 1:25 ratio of trypsin to total protein). Peptides were sequentially eluted from the column (first with 50 mM TEAB; then 0.2% formic acid (FA); and lastly, 50% acetonitrile (ACN)), with the flow-through pooled after each centrifugation. The peptides were concentrated via SpeedVac vacuum concentrator and frozen at -80 °C.

### SCX fractionation for building peptide LC-MS/MS library

150 µg of peptides were reconstituted in 0.5% FA by vortexing and sonication. Strong cation exchange (SCX) stage tips (Empore Cation Exchange-SR, Supelco Analytical) were activated with 100% ACN and then washed with 20% ACN, 0.5% FA. Peptide samples were loaded onto the tips and were washed with 20% ACN, 0.5% FA. Peptides were then separated into 8 fractions via elution with different concentrations of ammonium acetate (AA) (fractions 1-7 contained 20% ACN, 0.5% FA and either 45, 70, 135, 167, 200, 250 or 300 mM of AA; fraction 8 contained 5% ammonium hydroxide, 80% ACN) (pH 2.5 - 5.5). Each elution fraction was concentrated by SpeedVac vacuum concentrator and then frozen at -80 °C.

### Peptide Desalting

Each dried sample peptide at 50 µg and each dried SCX peptide fraction were reconstituted in 0.1% trifluoroacetic acid (TFA) and 2% ACN by vortexing and sonication. De-salting stage tips were prepared with two plugs of 3M™ Empore™ SDB-XC Extraction Disks (Fisher Scientific), which were activated with 100% ACN and washed with 0.1% TFA, 2% ACN (Kulak et al., 2014). The peptide sample was loaded onto the column, where peptides were washed with 0.1% TFA, 2% ACN before elution with 0.1% TFA, 80% ACN. Samples were then concentrated by speed-vacuum and then frozen at -80 °C.

### LC-MS

Peptides were reconstituted in 0.1% TFA, 2% ACN with vortexing and a 10-min sonication. These were centrifuged at max speed for 10 mins to pellet any impurities and the supernatant was used for LC-MS analysis.

The Nano-LC system, Ultimate 3000 RSLC (Thermo Fisher Scientific, San Jose, CA, USA) was set up with an Acclaim Pepmap RSLC analytical column (C18, 100 Å, 75 μm × 50 cm, Thermo Fisher Scientific, San Jose, CA, USA) and Acclaim Pepmap nano-trap column (75 μm × 2 cm, C18, 100 Å) and controlled at a temperature of 50 °C. Solvent A was 0.1% (v/v) FA and 5% (v/v) DMSO in water and solvent B was 0.1% (v/v) FA and 5% DMSO in ACN. The trap column was loaded with tryptic peptide at an isocratic flow of 3% ACN containing 0.05% TFA at 6 µL/min for 6 mins, followed by the switch of the trap column as parallel to the analytical column. The gradient settings for the LC runs, at a flow rate 300 nL/min, were as follows: solvent B 3% to 4% in 1 min, 4% to 25% in 75 min, 25% to 40% in 4 min, 40% to 80% in 1 min, maintained at 80% for 3 mins before dropping to 3% in 0.1 min and equilibration at 3% solvent B for 4.9 min.

Data-dependent acquisition (DDA) was carried out on SCX fractionated peptides to generate a reference spectral library using an Eclipse Orbitrap mass spectrometer (Thermo Fisher Scientific, San Jose, CA, USA). MS experiments were executed at positive mode with settings of nano electrospray ionization (nESI) spray voltage, S-lens RF, and capillary temperature level of 1.9 kV, 30%, 275 °C, respectively. The mass spectrometry data was acquired with a 3-s cycle time for one full scan MS spectra and as many data dependent higher-energy C-trap dissociation (HCD)-MS/MS spectra as possible. Full scan MS spectra feature ions at m/z of 375-1500, a maximum ion trapping time of 50 msec, an auto gain control target value of 4e5, and a resolution of 120,000 at m/z 200. An m/z isolation window of 1.6, an auto gain control target value of 5e4, a 30% normalized collision energy, a maximum ion trapping time of 22 ms, and a resolution of 15,000 at m/z 200 were used to perform data dependent HCD-MS/MS of precursor ions (charge states from +2 to +7). Dynamic exclusion of 30 sec was enabled.

Data-independent acquisition (DIA) was carried out on drug treated and SILAC samples using an Eclipse Orbitrap mass spectrometer (Thermo Fisher Scientific, San Jose, CA, USA) as above, with the following differences. A survey scan with a m/z range of 350 to 1400, a resolution of 120,000 at m/z 200, a normalized automatic gain control (AGC) target of 250%, a maximum ion trapping time of 50 ms was performed before 49 DIA windows with an m/z isolation window of 13.7, a precursor ion m/z range of 361-1033, a MS/MS scan range of m/z 200-2000, a resolution of 30,000 at m/z 200, an AGC of 1e6, a maximum ion trapping time of 55 ms and a normalized collision energy (NCE) of 30%.

DDA data for the SCX fractionated peptides were analysed with MaxQuant while DIA data were analysed with Spectronaut 16 using the direct-DIA (hybrid approach) with data from the SCX peptide fractions used as the extended library to improve the rate of peptide identification. Digestion enzyme specificity was set to trypsin and up to 2 missed cleavages were allowed. Mass tolerance for full scan MS and MS/MS scans were set at dynamic mode. Search criteria included carbamidomethylation of cysteine as a fixed modification, with oxidation of methionine and protein N-terminal acetylation as variable modifications. For SILAC labelled samples ¹³C₆¹⁵N₁ heavy isoleucine was selected for labeling in channel 2. Data were searched against UniProt human and *Plasmodium falciparum* (3D7) sequence database (December 2020). The false discovery rate (FDR) was set to 1% at peptide spectrum match (PSM), peptide and protein level. MaxLFQ of the intensity at MS/MS level was enabled for label free quantification and no imputation was enabled.

The software Perseus (version 2.0.7.0) was used for downstream bioinformatics analysis of the DIA data (Tyanova et al., 2016). Spectronaut output files were imported into Perseus where data was grouped by condition (drug treatment and incubation time). The MS2 Quantity was log₂-transformed. Missing values were replaced with random values within a window made to be consistent with the instruments limit of detection (width of 0.3 and down shift set at 1.8). To maximise possible hits, no filters were used for number of peptides per protein detected, number of samples a given protein was detected in across replicates or percent coverage of each protein for analysis of steady state data, however for SILAC data, unlabelled peptides must be detected in at least 75% of replicates for at least one treatment to be considered.

Statistical differences between groups were determined using a two-sample two-sided t-test with an FDR of 0.05. Nuclear encoded apicoplast-targeted and mitochondrial-targeted genes were annotated using published lists of predicted proteins (Boucher et al., 2018; van Esveld et al., 2021).

### Visualisation of mitochondrial membrane potential by fluorescence microscopy

Synchronised trophozoite-stage parasites were treated with DMSO, 1 or 50 µM doxycycline, or 50 µM carbonyl cyanide 3-chlorophenylhydrazone (CCCP, Sigma Aldrich, C2759) for 2.5 hrs at 37 °C. 25 nM MitoTracker™ Red CMXRos (Invitrogen) was then spiked into the cultures and parasites incubated for a further 30 mins while protected from light. Samples were fixed to glass coverslips, stained with 300 nM DAPI, and mounted onto glass slides for imaging. Data were processed using ImageJ (Version 1.53j) (Schindelin et al., 2012).

### Seahorse XFe96 extracellular flux analysis in *Plasmodium falciparum*

The oxygen consumption rate (OCR) of digitonin-permeabilised *P. falciparum* parasites was measured following a modified Seahorse XFe96 extracellular flux analyser assay protocol (Ramesh et al., 2023). Synchronised late-ring stage cultures were treated with DMSO or 25 µM doxycycline for ∼ 15 hrs, after which trophozoite stage parasites were isolated by magnetic purification. Parasite cultures were passed through a CS column (Miltenyi Biotec) held in a magnetic apparatus with a flow rate of ∼ 1 mL/min. The enriched iRBCs were eluted in CCM by removing the column from the magnetic apparatus. The iRBCs were treated with 0.05% (w/v) saponin, and the whole parasites collected by centrifugation. Parasites were washed in PBS and then resuspended in MAS buffer (220 mM mannitol, 70 mM sucrose, 10 mM KH₂PO₄, 5 mM MgCl₂, 0.2% (w/v) fatty acid-free bovine serum albumin (BSA), 1 mM EGTA and 2 mM HEPES-KOH, pH 7.4) supplemented with 25 mM malate and 0.002% (w/v) digitonin (Sigma) at 5.8 x 10⁷ parasites/mL. Parasites were seeded in a XFe96 cell culture plate pre-coated with Cell-Tak™ Cell and Tissue Adhesive (Corning 354240) at 5.8 x 10⁶ parasites per well, and the parasites were adhered to the bottom of the plate by centrifugation. The total volume of each well was adjusted to 175 µL with supplemented MAS.

The ports of the sensor cartridge (pre-hydrated the previous day with XF Calibrant Solution, stored overnight at 37 °C) were loaded with 25 µL of compounds and injected into the wells at set times during the assay. Atovaquone or doxycycline was injected from Port A to a final concentration of 5 µM and 25 µM respectively. N,N,N′,N′ - tetramethyl-p-phenylenediamine dihydrochloride (TMPD; Sigma-Alrich) and ascorbic acid were injected from Port B to a final concentration of 0.2 mM and 2 mM respectively and from Port C NaN₃ was injected to a final concentration of 10 mM. Measurements were taken for 2.5 mins after 20 seconds of mixing and 1 min of waiting. The baseline OCR had 5 measurements taken, 8 measurements were taken after the injection of Port A, and 5 measurements were taken after injections from both Port B and C. The OCR was calculated by subtracting the OCR of the background no-parasite wells that contained supplemented MAS, with 2-5 technical replicates included for each drug pre-treatment. The basal mOCR was calculated by subtracting the lowest OCR measurement following Port A injection from the final baseline OCR measurement. Spare respiratory capacity was measured by subtracting the basal mOCR from the highest measurement following Port B injection. Complex IV activity was determined by subtracting the lowest OCR measurement following Port C injection from the highest measurement following Port B injection.

### *Toxoplasma gondii* culture and Seahorse assays

*T. gondii* Δ*ku*80/TATi tachyzoites were cultured as previously described (Sheiner et al., 2011; Striepen & Soldati, 2007). Parasites were grown in a human foreskin fibroblast (HFF) primary cell line 53 (sourced from: ATCC, CRC1041; or the Peter MacCallum Cancer Centre) and DMEM (with 1% (v/v) foetal bovine serum and antibiotics) at 37 °C and 5% CO₂.

Experiments measuring the OCR of digitonin-permeabilised *T. gondii* parasites were conducted as previously described (Hayward et al., 2022; Hayward et al., 2021). Briefly, parasites were grown in the presence of either DMSO, 24 µM doxycycline, or 10 µM clindamycin for 1–2 days, egressed from host cells using a 26-gauge needle, and filtered through a 3 µM filter to remove host cell debris. Parasites were resuspended in non-supplemented base medium to 1.5 x 10⁷ cells/mL and starved for 1 hr at 37 °C to deplete endogenous ETC substrates. 1.5 x 10⁶ parasites were added to each well of a Seahorse XFe96 plate, pre-coated with Cell-Tak™ Cell and Tissue Adhesive and the parasites were adhered by centrifugation. Base medium was removed and replaced with 175 µL MAS buffer containing 0.002% (w/v) digitonin. Compounds were loaded into the sensor cartridge ports and injected into wells at set times during the assay. Carbonyl cyanide 4-(trifluoromethoxy)phenylhydrazone (FCCP) and malate were injected from Port A to final concentrations of 1 µM and 10 mM respectively. Atovaquone was injected from Port B to a final concentration of 1 µM, and TMPD and ascorbic acid were injected from Port C to final concentrations of 0.2 mM and 3.3 mM respectively. OCR was measured every 3 minutes three times before the first injection and three times after each injection. Malate- and TMPD-elicited mOCR were calculated by subtracting the non-mitochondrial OCR (the values following atovaquone injection) from the OCR value obtained after malate and TMPD injections respectively.

### Analysis of Seahorse Data

Data were exported from the Seahorse Wave Desktop software (Agilent Technologies). A linear mixed effects model was applied to the data, as previously described (Hayward et al., 2022): the error between plates (biological replicates) and wells (technical replicates) were set as random effects, and the mOCR values between cell lines and days on drug as fixed effects. Analysis of the least square means of the values was performed in the R software environment (Hayward et al., 2022). Statistical differences between those values were tested through ANOVA (linear mixed effects) with a post hoc Tukey test.

### Native PAGE and ETC complex III, IV and V detection

Native PAGE experiments to detect complex IV and V were performed as previously described (Lacombe et al., 2019; Shikha et al., 2022) with minor modifications. Briefly, RHΔ*ku*80 TATi tachyzoites were cultured with 1, 25 or 50 µM doxycycline or 0.002 or 10 µM clindamycin for 24 hrs. For in-gel complex IV enzymatic activity detection, 1 x 10^7^ parasites were resuspended in solubilisation buffer (50 mM NaCl, 50 mM imidazole, 2 mM 6-aminohexanoic acid, 1 mM EDTA–HCl pH 7.0, 2% (w/v) n-dodecylmaltoside). Samples were incubated on ice and then solubilised membrane proteins isolated by centrifugation at 16,000 × g at 4 °C for 15 mins. Samples were combined with glycerol and ponceau S to a final concentration of 6.25% and 0.125% respectively and then separated on a 4–16% Bis-Tris gel. NativeMark was used as a molecular weight marker. To detect complex IV activity, gels were incubated in a 50 mM KH2PO4, pH 7.2, 1 mg ml^−1^ cytochrome *c*, 0.1% (w/v) 3,3′-diaminobenzidine tetrahydrochloride solution. For complex V detection 5 x 10^6^ parasites were resuspended in nativePAGE sample buffer (Invitrogen) containing 1% (w/v) n-dodecylmaltoside. Samples were incubated on ice and then solubilised membrane proteins isolated by centrifugation at 16,000 × g at 4 °C for 15 mins. Samples were combined with sample buffer containing Coomassie G250 (NativePAGE) and proteins separated on a NativePAGE 3–12% Bis-Tris gel. NativeMark was used as a molecular weight marker. Proteins were then transferred to polyvinylidene fluoride (PVDF) membrane and immunolabelling performed using anti-ATPβ antibody (Agrisera AS05085, 1:5000).

To generate a *T. gondii* line in which we could simultaneously detect complexes III and IV, we incorporated a HA epitope tag into the *Tg*Cox2a locus of an existing parasite line in which *Tg*QCR11 was FLAG-tagged (Hayward et al., 2021). We amplified a 3×HA tag using the primers Cox2a 3’rep fwd and rvs (described previously, (Seidi et al., 2018)), which contain 50 bp homologous flanks either side of the *Tg*Cox2a stop codon, and a gBlock encoding a 3×HA tag (IDT) as template (also described previously, (Hayward et al., 2021)). We mixed the PCR product with an existing vector termed *Tg*Cox2a 3’ sgRNA in pSAG1::Cas9-U6 (described previously, (Seidi et al., 2018)), which encodes a single guide RNA targeting the 3’ region of the *Tg*Cox2a open reading frame and a Cas9-GFP protein for CRISPR-based genome editing. We transfected the DNA mix into *Tg*QCR11-FLAG parasites and cloned GFP-positive parasites two days following transfection using flow cytometry. We verified successful integration of the HA tag into the *Tg*Cox2a locus through PCR screening, using the primers Cox2a 3’ screen fwd and rvs (**Supp Figure 5**; primers described previously (Seidi et al., 2018)).

To test the effects of doxycycline and clindamycin treatment on complex III and IV abundance, we cultured parasites in the presence of DMSO, 25 µM doxycycline or 10 µM clindamycin for one or two days. For BN-PAGE, we solubilised parasite proteins in NativePAGE sample buffer (Thermo Fisher Scientific) containing 1% (v/v) Triton X-100, 2 mM EDTA and 1× cOmplete protease inhibitor cocktail (Merck). We separated 2.5 × 10^6^ parasite equivalents on a precast 4-16% Bis-Tris Native PAGE gel (Thermo Fisher Scientific), transferred proteins to a PVDF membrane, and probed by western blotting. For SDS-PAGE, we solubilised proteins in LDS sample buffer (Thermo Fisher Scientific) containing 2.5% (v/v) b-mercaptoethanol. We separated 2.5 × 10^6^ parasite equivalents on a precast 12% Bis-Tris NuPAGE gel (Thermo Fisher Scientific), transferred proteins to a nitrocellulose membrane, and probed by western blotting. All membranes were blocked in Tris-buffered saline containing 4% (w/v) skim milk powder. For the western blotting, membranes were probed with rat anti-HA (1:2,000 dilution; Merck clone 3F10, catalogue number 11867423001), mouse anti-FLAG (1:1,500 dilution; Merck clone M2, catalogue number F3165), or rabbit anti-*Tg*Tom40 (1:2,000 dilution, (van Dooren et al., 2016)) primary antibodies, and horseradish peroxidase-conjugated goat anti-rat (1:10,000 dilution; Abcam catalogue number ab97057), goat anti-mouse (1:10,000 dilution; Abcam catalogue number ab6789) or goat anti-rabbit (1:10,000 dilution; Abcam catalogue number ab97051) secondary antibodies. Blots were imaged with a BioRad ChemiDoc MP imaging system.

## Acknowledgment

LS is funded by Wellcome Investigator Award (217173_Z_19_Z). We thank Glasgow honours students Ciara Loughrey and Dzachery Zainudden for helping with the in-gel CIV assays in *Toxoplasma*. This research was supported by an Australian National Health and Medical Research Council (NHMRC) Project grant (GNT1140906) Investigator Fellowship (GNT2009732) to DAS and an NHMRC Ideas grant (GNT1182369) to GGvD. We thank the Mito Foundation for the provision of instrumentation through research equipment grants to DAS and a scholarship to LM-W. We thank the Bio21 Mass Spectrometry and Proteomics Facility (MMSPF) for the provision of instrumentation, training, and technical support. The mass spectrometry proteomics data have been deposited to the ProteomeXchange Consortium via the PRIDE (Perez-Riverol et al., 2024) partner repository with the dataset identifier PXD071325.

---

Supplementary Tables 1-7 in Excel file.

**Supp Figure 1.**
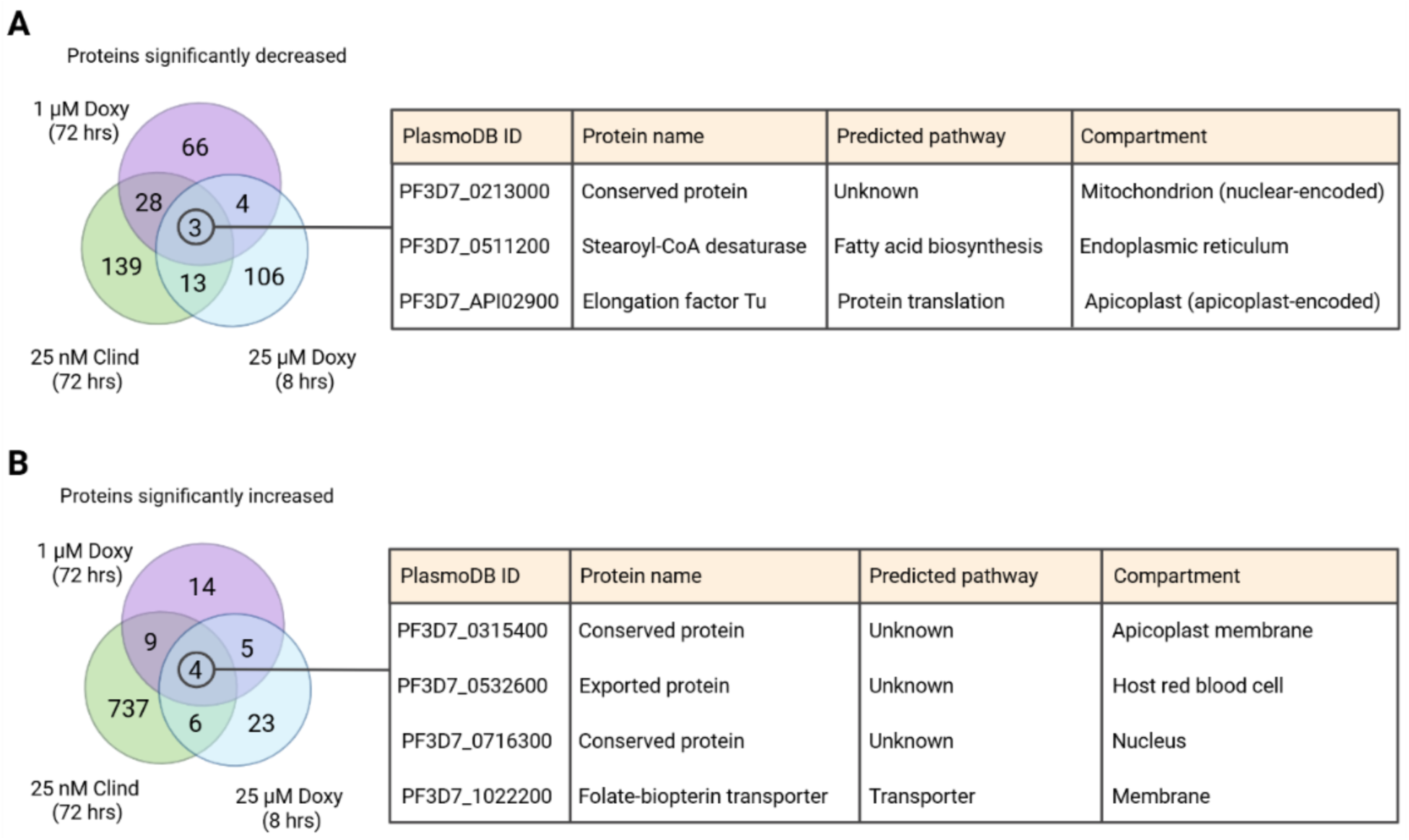
Common proteins significantly A) decreased or B) increased in delayed death (72 hrs with 1 µM doxycycline or 25 nM clindamycin) or high doxycycline (25 µM) treatments. p value < 0.05, fold change > 1.5 x.

**Supp Figure 2.**
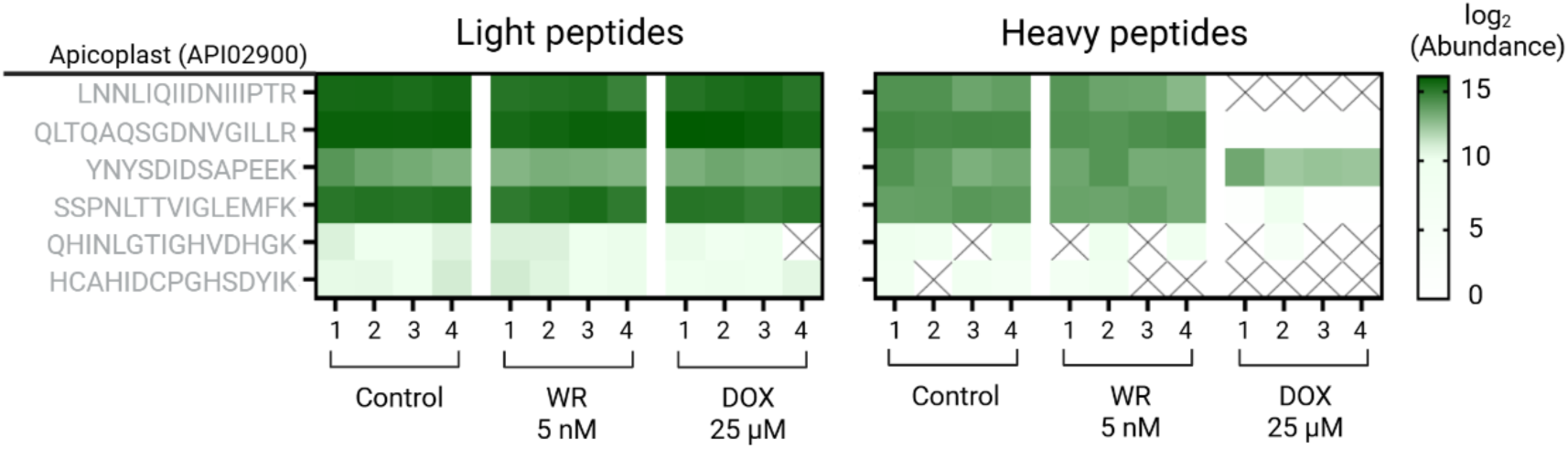
Inhibition of apicoplast translation occurs with doxycycline but not WR treatment in *P. falciparum*. Parasites were treated with either DMSO, WR99210 (WR, 5 nM) or doxycycline (DOX, 25 µM) for 8 hrs. After 3 hrs of treatment, media was exchanged from that containing light to heavy isoleucine. Heat map showing the log₂ abundance of the light and heavy isoleucine-containing peptides of organellar-encoded proteins. The four replicates for each treatment are shown. Only light peptides detected in at least 75% of replicates for at least one treatment are shown.

**Supp Figure 3.**
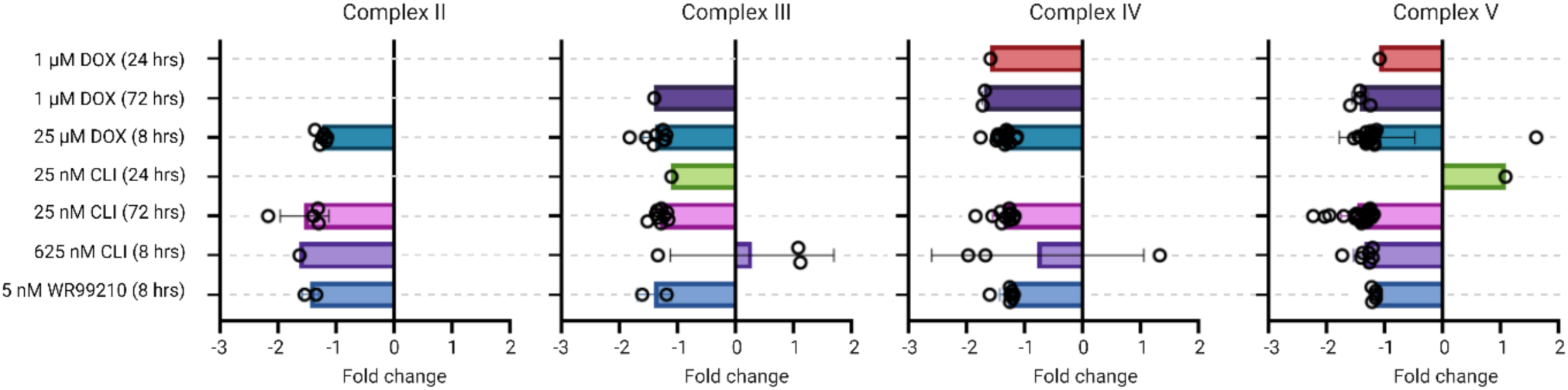
Drug treatments only have minor overall impacts on the steady state abundance of ETC subunits. Abundance of nuclear genome encoded subunits of complexes II-V of the electron transport chain was assessed by LC-MS. The fold change of all proteins with a p value < 0.05 are displayed, which show a trend towards a decrease across all complexes following drug exposure.

**Supp Figure 4.**
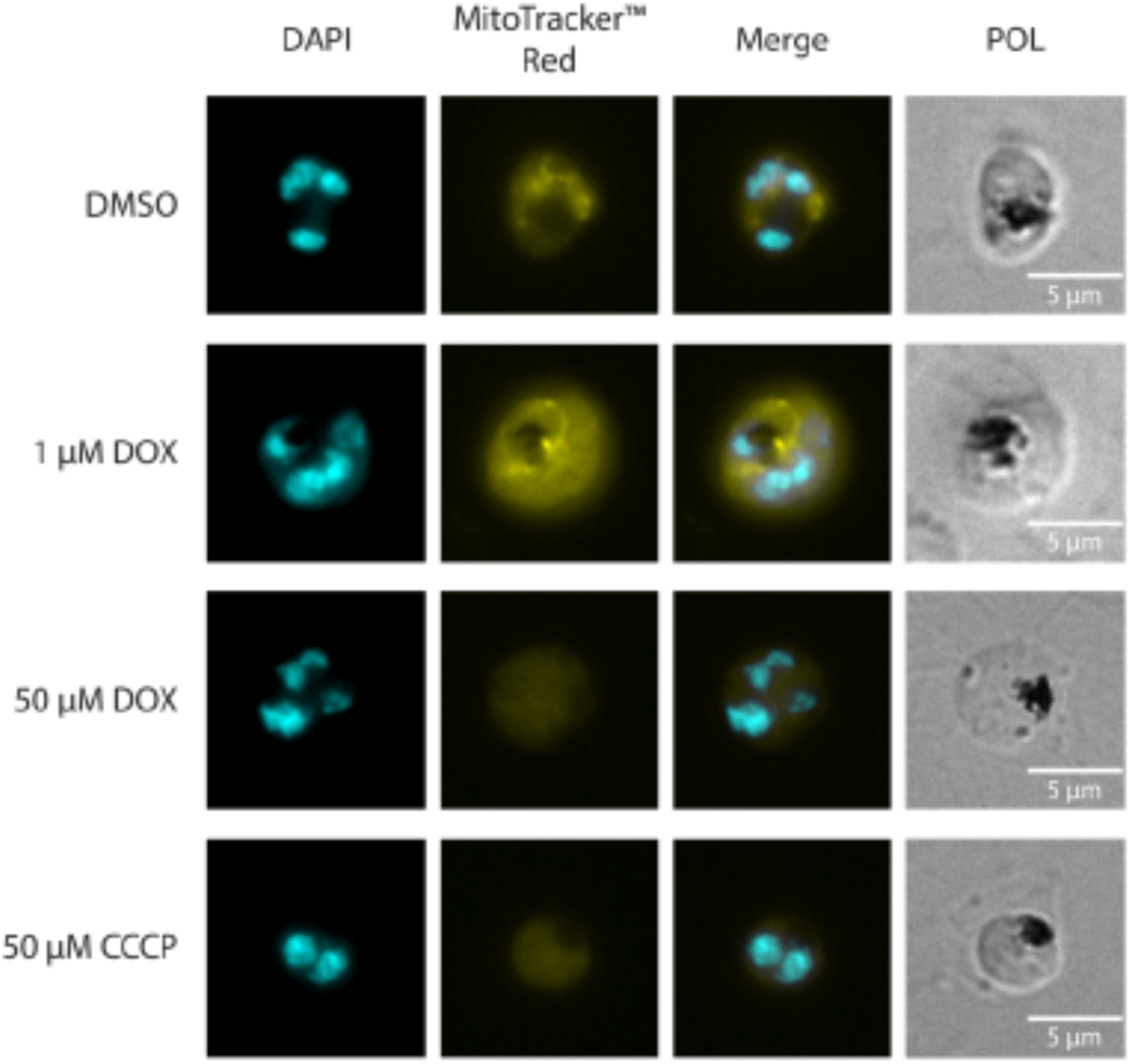
High doxycycline treatment results in depolarized mitochondrion. 3D7 *Plasmodium falciparum* trophozoite stage parasites were treated with DMSO, 1 or 50 µM doxycycline (DOX), or 50 µM of mitochondrion decoupler carbonyl cyanide 3-chlorophenylhydrazone (CCCP) for 3 hrs. Parasites were stained with 25 nM MitoTracker™ Red CMXRos (yellow) before samples were fixed with 3.7% (v/v) paraformaldehyde (PFA) and stained with 300 nM DAPI (cyan) for imaging by fluorescence microscopy. Representative data are presented (n = 1).

**Supp Figure 5.**
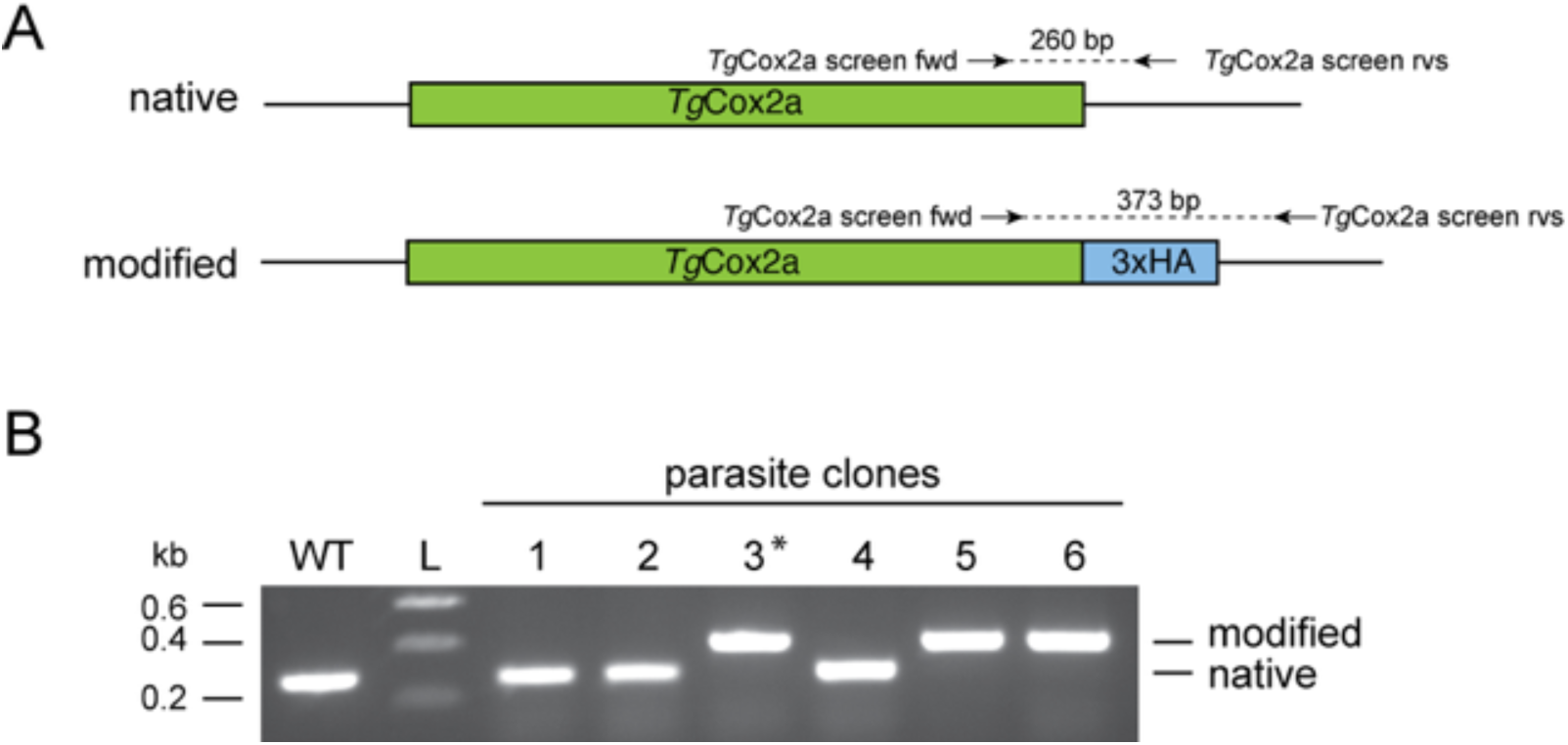
Generating a *T. gondii* parasite line for simultaneously detecting complexes III and IV. **A)** Schematic depicting the *Tg*Cox2a locus before and after integration of a 3×HA epitope tag into the 3’ region of the open reading frame of the gene. The approximate position of the primers used for subsequent PCR analysis and the expected sizes of the PCR products in the native and genetically modified loci are indicated. **B**) PCR screen of parasite clones testing for integration of a 3×HA tag into the *Tg*Cox2a locus using genomic DNA from parasite clones as a template, and genomic DNA from wild type (WT) parasites as a control for the expected size of the native locus. The asterisk denotes the clone selected for subsequent experiments.

## Notes

### Competing Interest Statement

The authors have declared no competing interest.

### Summary of Updates

Added references, revised description of seahorse assay to avoid potential confusion

